# Assessment of the Human Placental Microbiome in Early Pregnancy

**DOI:** 10.1101/2022.07.04.498741

**Authors:** Vassilena Sharlandjieva, Alexander G. Beristain, Jefferson Terry

**Affiliations:** BC Children’s Hospital Research Institute, Vancouver, BC V5Z 4H4; BC Children’s Hospital, Vancouver, BC V6H 3V4

**Author notes:** Corresponding author Jefferson Terry, Clinical Associate Professor, Department of Pathology and Laboratory Medicine, University of British Columbia, BC Children’s Hospital, 4480 Oak Street, Room 2H60, Vancouver, BC V6H 3V4.

## Abstract

Bacteria derived from the maternal circulation have been suggested to seed the human placenta during development leading to an intrinsic placental microbiome. This concept has become controversial as numerous studies suggest that the apparent placental microbiome is mostly, if not completely, comprised of contaminants. If the maternal circulation seeds the placenta then there should be an increase in abundance and diversity of detectable bacteria with onset of maternal perfusion of the placenta around 10 weeks gestational age; however, if only contaminants are present then there should be no significant evolution of the placental microbiome with increasing gestational age. This pilot study addresses whether bacterial abundance and diversity increases in human placenta and whether there is an associated shift in the immunophenotype of the decidual immune cell complement before and after initiation of placental perfusion. Human placental and decidual tissue from 5-19 weeks gestational age are assessed by quantitative 16S polymerase chain reaction (PCR), 16S gene sequencing, and immunological flow cytometry studies. A weak positive correlation between placental bacterial abundance and gestational age is identified but is not statistically significant. No significant changes in bacterial diversity are found with increasing gestational age. The proportion of decidual activated memory T helper cells increases with gestational age but no change was observed in other lymphocyte subsets. This pilot study does not strongly support bacterial colonization of the placenta after initiation of maternal perfusion; however, the minor trends towards increases in bacterial abundance and activated memory T helper cells may represent an early stage of this process. Additional investigations in larger cohorts are warranted.

## Introduction

Bacteria appearing to be derived from the maternal oral cavity and intestinal lumen have been reported in placental tissue, amniotic fluid, and meconium from preterm and term pregnancies [1-7]. These reports, which assess for the presence of bacteria through culture and molecular-based methodologies, imply that transiently circulating bacteria from the maternal oral cavity and/or intestine seed the placenta *in utero* via the maternal bloodstream. Data demonstrating an intrauterine placental microbiome and translocation of bacteria from the placenta to the fetus support the prenatal placental bacterial colonization hypothesis [4,8-13]. Other studies, however, have shown that bacteria or bacterial DNA detected in placental and fetal tissues are most likely contaminants [14-24], raising doubt that a true placental microbiota exists and proposing that if bacterial seeding of the placenta and fetus does occur, it is likely around the time of delivery.

If a placental microbiome originates from maternal circulation-derived bacteria, then an increase in bacterial abundance and diversity should become detectable at the onset of placental perfusion by the maternal circulation. After conception, the developing placenta and embryo are sequestered from the maternal circulation until approximately 10 weeks gestational age, when the maternal blood space of the developing placenta connects to maternal vessels in the adjacent decidua and exposes the placenta directly to maternal blood. After this point in gestation the abundance and diversity of bacteria detectable in the placenta would be expected to increase with increasing gestational age and cumulative exposure to the maternal bloodstream. The maternal decidua, which is consistently perfused during this timeframe, should maintain a relatively consistent microbiome signature. An increase in placental bacterial abundance and/or diversity relative to the maternal decidua after initiation of perfusion would be consistent with early intrauterine development of a placental microbiome.

Another potential indicator of the establishment of a placental microbiome is the reaction of the resident maternal immune system in the adjacent decidual tissue. Decidual immune cells, which are present at the maternal-fetal interface, may play a permissive role in placental microbiota development. The oral and gut microbiota can affect the differentiation and phenotypic characteristics of mucosal leukocytes [8,25,26], suggesting that the establishment of microbiota at the maternal-fetal interface may be associated with similar detectable shifts in decidual immune cells.

This pilot study examines the presence, timing, and composition of bacteria at the fetal-maternal interface, using quantitative 16S gene PCR and 16S gene sequencing, and the decidual immune cell reaction by flow cytometric analysis of the complement of decidual T cells to address the hypothesis that development of an intrauterine placental microbiome is associated with an increase in bacterial abundance and diversity in placental tissue and changes in resident decidual T cells after onset of maternal perfusion in the late first trimester of pregnancy.

## Materials and Methods

Quantitative 16S rRNA qPCR and 16S gene sequencing measures the abundance and composition of bacteria at the fetal-maternal interface. Decidua, which is perfused throughout pregnancy and will acquire the same contaminants as the placental samples, is employed as an internal control and comparator for contamination and microbiome development respectively; if a placental microbiome develops after onset of perfusion, this will be detectable as an increase in abundance and/or diversity relative to decidua (Supplemental Fig. 1). Flow cytometric analyses of the complement of decidual T cells during before, during, and after the onset of perfusion address the hypothesis that development of an intrauterine placental microbiome is associated with changes in resident decidual T cells.

### Study Cohort

Ethics approval is obtained from the University of British Columbia Research Ethics Board (H17-03582). 25 women presenting for pregnancy termination at BC Women’s Hospital that meet eligibility criteria and provide written informed consent to participate are enrolled. The consenting process and all experiments are performed in accordance with applicable guidelines and regulations. Criteria for inclusion are gestational age 5-19 weeks, elective termination, and termination by dilatation and evacuation or curettage performed with sterile equipment and aseptic technique. Exclusion criteria are history at presentation of a chronic maternal inflammatory condition, acute maternal inflammatory condition within 2 weeks of the termination procedure, history of fetal, placental, or other obstetrical abnormality, invasive instrumentation during the present pregnancy prior to termination, assisted conception in the present pregnancy, history of inherited disease or recurrent pregnancy loss, and antibiotic use at the time of presentation. The study cohort is described in Supplementary Table S1.

### Tissue Samples

Sampled tissues include placental villi and maternal decidua. Organ tissues are also sampled if present. Tissue samples are collected as participants are enrolled and samples become available. A single operator handles all samples with aseptic technique using sterile equipment in a laminar flow hood. Samples are washed with sterile phosphate-buffered saline (PBS) to remove maternal blood and reduce surface contamination. Samples of placenta, decidua, and fetal organ tissues identifiable by gross examination are taken. Tissue and PBS wash samples are frozen within 6 hours of retrieval and stored at −80 °C prior to analysis. Representative portions of each tissue sample are formalin fixed and processed for histological assessment using standard protocols and reagents to confirm tissue type, to assess for abnormal inflammation based on established histological criteria [27,28], and to assess for presence of bacteria. If abnormal inflammation is identified in any sample, all samples from that participant are excluded from further analysis.

### Microbiome Analysis

Microbiome analysis is performed by Microbiome Insights (Vancouver, BC, Canada). DNA is extracted from 0.25 g of tissue, PBS wash, and PBS negative control samples using a PowerMag Soil DNA Isolation Bead Plate (MPBio, Irvine, CA, USA) following the manufacturer instructions with a KingFisher robot (ThermoFisher, Waltham, MA, USA). Quantitative 16S ribosomal ribonucleic acid (rRNA) gene PCR is performed using bacteria-specific forward (300 nM 27F, 5’-AGAGTTTGATCCTGGCTCAG-3’) and reverse (300 nM 519R, 5’-ATTACCGCGGCTGCTGG-3’) primers. 25 µl reactions using iQ SYBR Green Supermix (Bio**-**Rad, Hercules, CA, USA) are run, in triplicate, on a StepOne Plus instrument (Applied Biosystems, Waltham, MA, USA) using the following protocol: initial denaturation at 95 °C for 3 minutes followed by 40 cycles of denaturation at 95 °C for 20 seconds, annealing at 55 °C for 30 seconds, and extension at 72 °C for 30 seconds. Full-length bacterial 16S rRNA gene cloned into the pCR4-TOPO plasmid is used as an amplification standard. 16S copy number per sample is determined by linear regression based on the cycle threshold (CT) values of 10-fold serial dilutions of the bacterial amplification standard. Template-free PBS is included as a negative control and yields no detectable amplification product. The mean CT and 16S copy quantification values for each sample are included in Supplementary Table S2.

DNA extracted for quantitative 16S PCR for all samples except negative PBS control, which yielded no amplifiable DNA, are used for 16S gene sequencing. Bacterial 16S rRNA genes are PCR-amplified with dual-barcoded primers targeting the V4 region (515F 5’-GTGCCAGCMGCCGCGGTAA-3’ and 806R 5’-GGACTACHVGGGTWTCTAAT-3’), as previously described [29]. Amplicons are sequenced on an Illumina MiSeq system (San Diego, CA, USA) using the 300-bp paired-end kit (v.3) yielding at least 5,000 quality paired-end 250-bp reads per sample. Sequencing quality for R1 and R2 is determined using FastQC 0.11.5. Data are analyzed using MOTHUR (v. 1.39.5) [30], including assembly of paired ends, quality filtering, binning of 97% identical operational taxonomic units (OTUs), and calculation of relative OTU abundance. Greengenes database (v. 13_8) is used for taxonomic identification. Reagent-based contaminants are controlled for by co-sequencing template-free PBS controls and extraction reagents. OTUs are considered reagent contaminants and removed from further analysis if their mean abundance in reagent controls (PBS and/or extraction reagents) is ≥ 25% of their mean abundance in the tissue samples. The FASTQ data generated and analyzed during this study are available in the SRA database under accession number PRJNA70705 (https://www.ncbi.nlm.nih.gov/sra/PRJNA707075).

### Decidual Immune Cell Phenotyping

Sampled decidual tissue is washed, minced, and enzymatically digested to produce a cell/tissue suspension as previously described [31]. Decidual leukocyte enrichment is performed by discontinuous Percoll density gradient centrifugation (layered 40%/80%) after which enriched decidual leukocytes are > 90% CD45+ cells as assayed by flow cytometry. Following washing in fluorescence-activated cell sorting buffer (PBS 1X, 1:1000 FVD520, 1% FBS), 1×106 total decidual leukocytes cells are incubated with FcBlock (ThermoFisher) for 10 minutes to block nonspecific binding. Cells are then incubated with different combinations of fluorescent-conjugated antibodies directed against specific extracellular markers for 30 minutes at 4°C. FVD520 is used to exclude dead cells. CD45 and CD3 markers define the T cell population, which is assessed for central and effector memory cells by CD45RO, CD3, CD4, CD8, CD25, and FoxP3 expression. Cells are analyzed on the BD LSRII flow cytometer (Franklin Lakes, NJ, USA) and data analyzed using FlowJo software (Tree Star, Inc., Ashland, OR, USA). Antibodies used are summarized in Supplementary Table S3.

### Statistical Analysis

Quantitative 16S PCR data is assessed by Welch’s t-test, Pearson correlation coefficient, and Kruskal-Wallis rank sum test. Differences are accepted as significant at *p* values less than .05. Microbiome alpha diversity is estimated by Shannon diversity index on raw OTU abundance tables after reagent contaminant filtering. The significance of diversity differences is tested by analysis of variance (ANOVA). To estimate beta diversity, Bray-Curtis indices are computed after excluding OTUs with a count of less than 3 in at least 10% of the samples. Beta diversity is visualized using Principal Coordinate Analysis (PCoA) ordination. Variation in community structure is assessed with permutational multivariate analyses of variance (PERMANOVA) using 9999 permutations for significance testing. Differentially abundant OTUs are assessed using DESeq2 [32]. All microbiome analyses are conducted in the R environment (version 3.5).

For flow cytometry data analysis, samples are grouped by gestational age into “early” (5 to 7 weeks gestation), “mid” (8-10 weeks) and “late” (over 10 weeks) to represent the pre-perfusion, onset of perfusion, and post-perfusion timeframes and the proportions of various cell types are compared between these groups. Flow cytometry data calculations are carried out using GraphPad Prism software (v. 8). For multiple comparisons, Tukey-Kramer tests are performed following two-way ANOVA. The differences are accepted as significant at *p* values less than .05. Non-parametric Spearman correlation was performed to test for correlation between placental OTUs and the proportion of activated CD4+ T cells, activated CD8+ T cells, memory Treg cells, and the ratio of Th1:Th2 cells.

## Results

### Study Cohort

The average participant age is 28 years (range 19-40), average gestational age at delivery is 9.5 weeks (range 5.3-17.1), and average pre-pregnancy body-mass index is 26.3 (range 18.1-38.8). A total of 24 placental, 22 decidual, and 6 organ tissue samples are obtained. The gestational age range of the placenta samples is 5.3-17.1 weeks, decidua 5.3-14.4 weeks, and organ tissue 11.0-17.1 weeks. Nine PBS wash samples are included to assess superficial bacteria/bacterial DNA removed by sample washing.

### Histological Analysis

All sampled placental, decidual, and organ tissues are confirmed as appropriately identified and free of other tissue types by histological assessment. No abnormal inflammation is identified in any tissue sample. Bacteria are not evident on hematoxylin and eosin staining in any of the sampled tissues.

### Quantitative 16S PCR

The alternative hypothesis of this study expects an increase in bacterial abundance in the placenta after initiation of maternal perfusion around 10 weeks gestational age. This is assessed by quantitative 16S PCR, which demonstrates the presence of amplifiable 16S DNA in all samples but no apparent increase in abundance with gestational age in the placental or decidual samples (Fig. 1). Amplification of template-free PBS demonstrated no amplifiable product after 40 cycles. Welch’s t-test is used to determine if there is a difference in the mean 16S copies in placental and decidual samples obtained before and after 10 weeks gestational age (pre-and post-perfusion). No significant difference is found for the placental (t = −0.25, two-tail *p* = .81) or decidual (t = 0.087, two-tail *p* = .93) samples, indicating no significant increase in bacteria, as measured by the presence of 16S DNA, after the onset of maternal perfusion.

**Figure 1.**
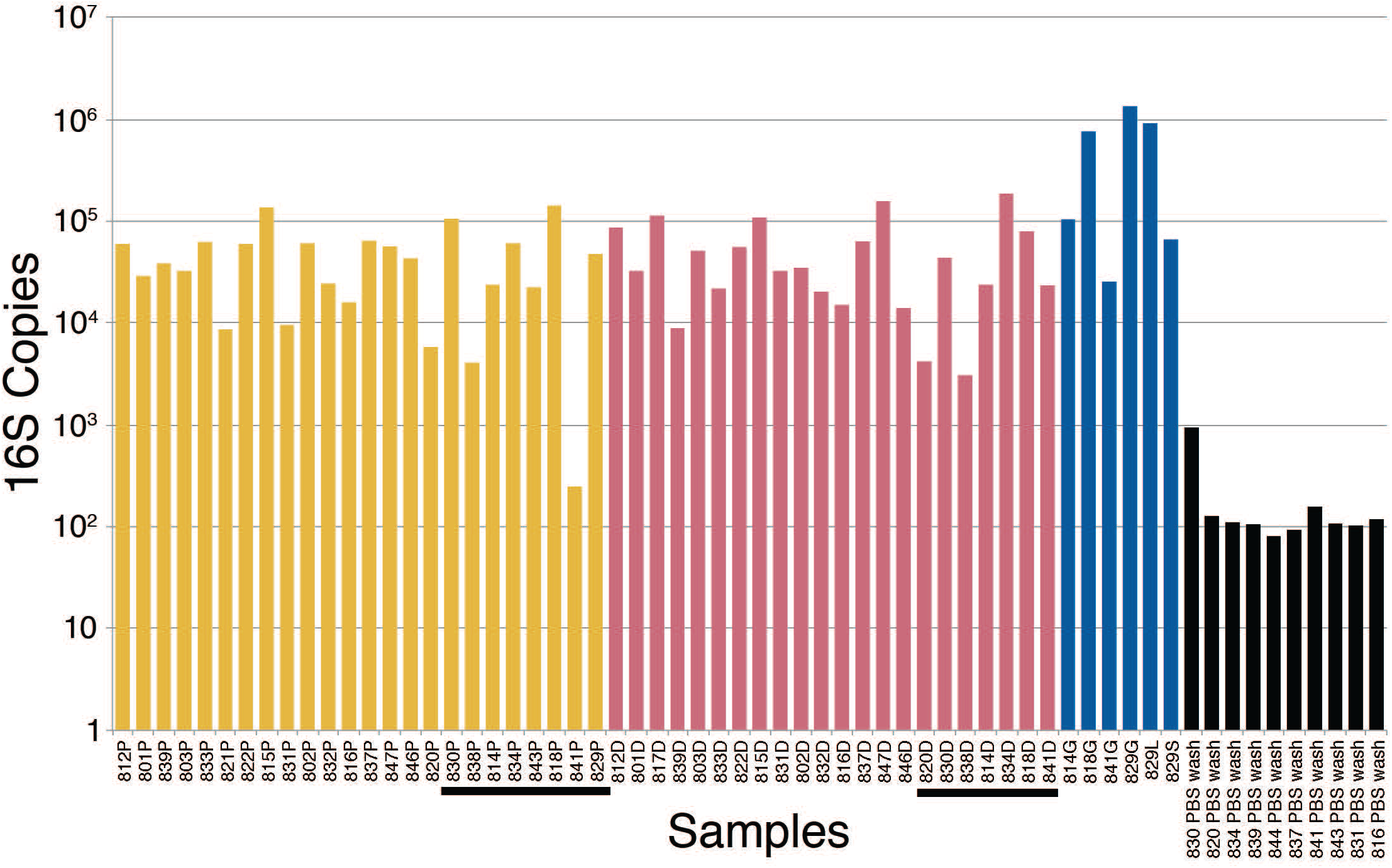
16S rRNA gene PCR bacterial quantification in placenta (yellow), decidua (red), fetal tissue (blue), and PBS wash (black) samples. The tissue samples are in order of increasing gestational age from left to right. Post perfusion (10 weeks gestational age and above) placental and decidual samples are underlined. P = placenta, D = decidua, G = gut (intestine) S = spleen, L = liver, PBS = phosphate buffered saline.

Computing Pearson correlation coefficients between the four sample sets (placenta, decidua, fetal tissue, PBS wash) and gestational age assesses correlation between 16S copy number and gestational age. There is a weak positive correlation between 16S copy number and gestational age for placenta (*r*(19) = .28, *p* = .22) and a moderate positive correlation in the organ tissue samples (*r*(5)= .47, *p* = .35), but neither are statistically significant. There is a no significant correlation between 16S copy number and decidua (*r*(20) = .05, *p* = .81) or PBS wash (*r*(8) = -.13, *p* = .72).

Kruskal-Wallis testing is performed to evaluate the differences in 16S copy number between the four groups of samples (placenta, decidua, fetal tissue, PBS wash), which demonstrates a significant difference (H(3) = 29.73, *p* = < .001). Follow-up Dunn testing with Benjamini-Hochberg correction for false discovery finds significant differences between placenta and organ tissue samples (*p =* .041) and decidua and organ tissue samples (*p =* .045), consistent with increased bacterial 16S DNA in the sampled fetal tissues compared to placenta and decidua (Fig. 1). The PBS wash samples showed significantly less 16S copies than the solid tissue samples (all *p* < .001), consistent with dilution during the PBS wash step. No significant difference in 16S copy number was observed between placenta and decidua (*p =* .86), indicating no significant increase in placenta bacteria/bacterial DNA relative to decidua after the onset of maternal perfusion.

### 16S Sequencing

All tissue and PBS wash samples are submitted for 16S gene sequencing and show amplification product (Supplemental Fig. S2). The per-base Q30 forward and reverse read quality (Phred) scores are good (Supplemental Fig. S3 and S4). After quality and reagent contaminant filtering and clustering reads with minimum 97% similarity, the dataset contains 7248 OTUs with an average of 6607 quality-filtered reads per sample. The number of quality reads obtained per sample by site is shown in Supplemental Fig. S5.

The alternate hypothesis of this study anticipates an increase in placental bacterial diversity after initiation of maternal perfusion around 10 weeks gestational age. Decidua, which is consistently exposed to the maternal circulation, would be expected to show relatively consistent bacterial diversity over the same timeframe. Thus the pre-perfusion placental samples are expected to show a relative lack of diversity compared to contemporaneous decidual samples and that should begin to equilibrate after 10 weeks. Compositional similarity (alpha diversity) for placenta and decidua <10 weeks and ≥10 weeks (samples 816, 820, and 821 are excluded due to a data processing error) is demonstrated in Fig. 2. There is no significant difference in the Shannon diversity index between <10 weeks and ≥10 weeks (ANOVA, *F*(3,19) = 1.366, *p* = .28). Microbiome similarity for placenta and decidua <10 weeks and ≥10 weeks (excluding samples 816, 820, and 821) (beta diversity) is calculated as Bray-Curtis dissimilarities for sample OTUs and expressed by principal coordinate analysis (Fig. 3); no statistically significant difference is identified (PERMANOVA, R2 = 0.1491, *p* = .315). Comparing diversity by site of sampling shows no significant difference in alpha diversity (ANOVA, *F*(5,56) = .80, *p* = .55; Supplemental Fig. 6a) or beta diversity (PERMANOVA, R2 = 0.0881, *p* = .418; Supplemental Fig. 6b). Differential abundance testing (DESeq2, R package) did not identify any taxonomic compositional differences between placenta, decidua, organ tissues, and PBS wash samples. These data demonstrate no significant difference in the bacterial diversity of placental or decidual samples before and after onset of maternal perfusion.

**Figure 2.**
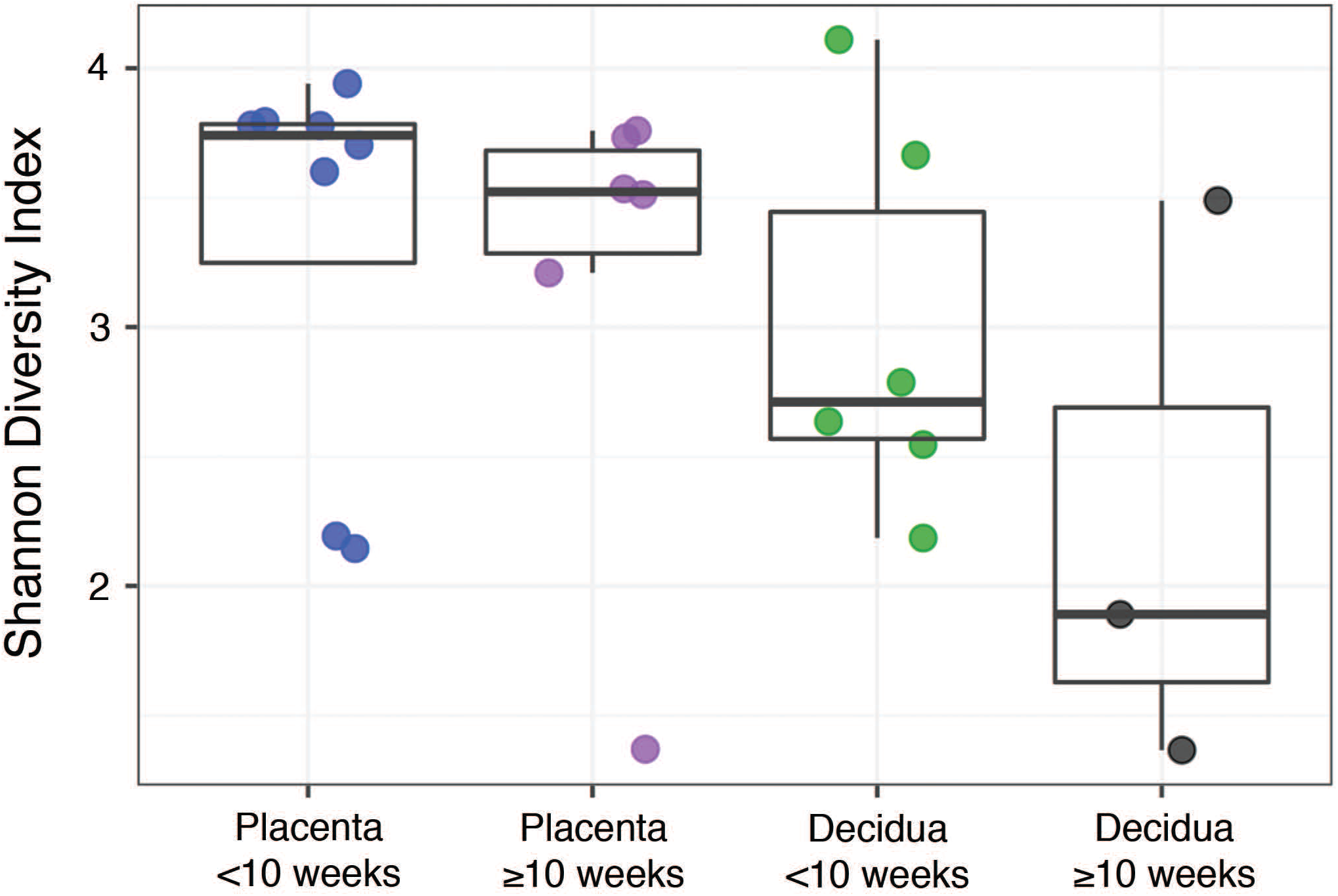
Alpha diversity between placental and decidual samples before and after 10 weeks gestational age.

**Figure 3.**
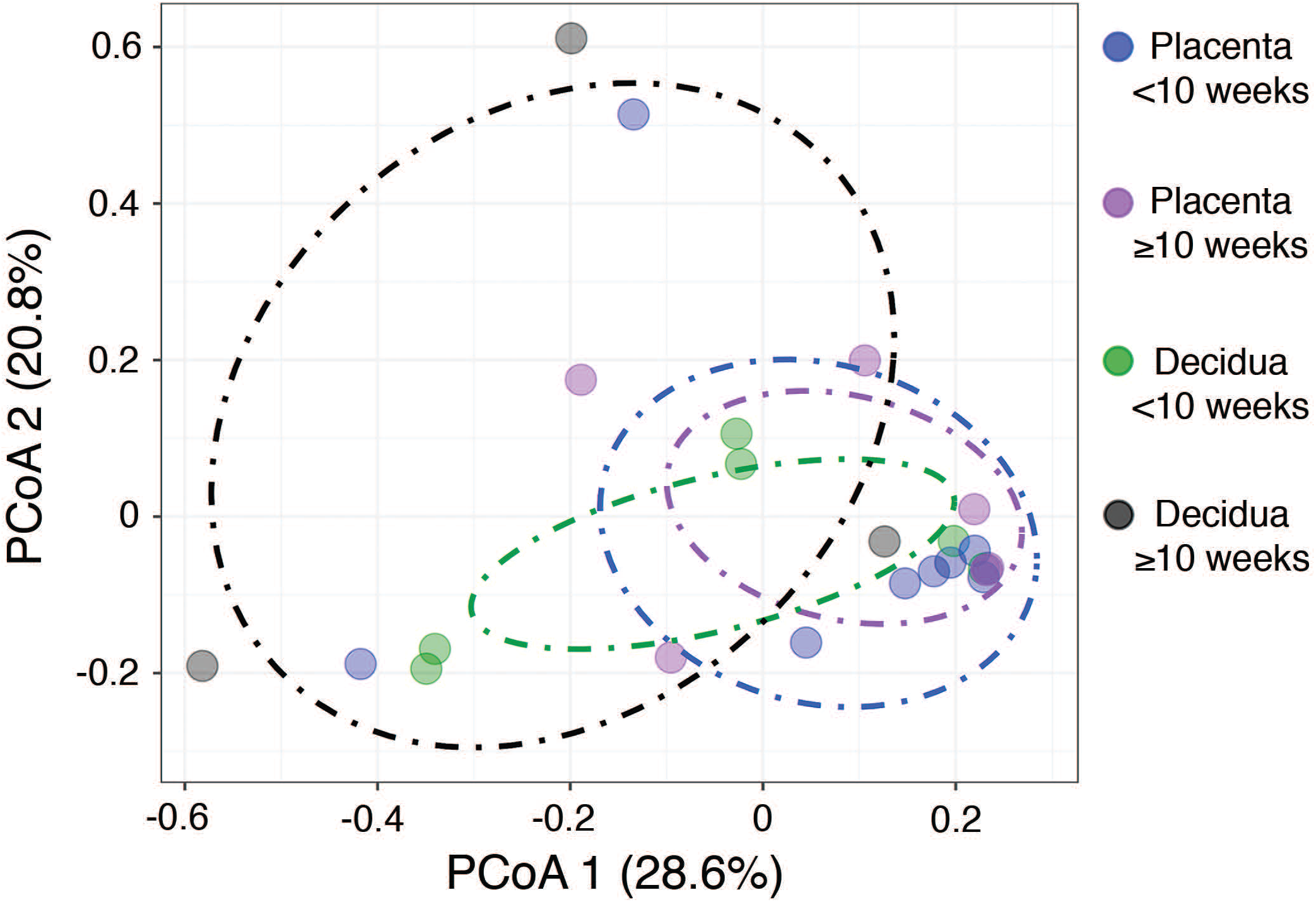
Beta diversity between placental and decidual samples before and after 10 weeks gestational age.

The taxonomic composition of the tissue samples is primarily comprised of human commensal bacteria found in the gastrointestinal tract and vagina (Fig. 4, Supplemental Fig. S7-S11). A subset of the decidual and placental samples shows increased relative abundance of Actinobacteria and Firmicutes (Supplemental Fig. S7). Assessment of these specimens at the genus/species level (Supplemental Fig. S11) shows increased levels of *Atopobium, Gardnerella*, and *Lactobacillus*, suggesting that the increased Actinobacteria and Firmicutes content of these samples are due to increased vaginal contamination. The decidual sample from specimen 846 showed increased Firmicutes that is primarily due to a single *Staphylococcus* species. *Chryseobacterium* and *Stentrophomonas*, which are typically environmental bacteria, are increased in a subset of samples. The PBS wash samples, which represent superficial bacteria present on the samples have a taxonomic composition generally similar to the tissue samples.

**Figure 4.**
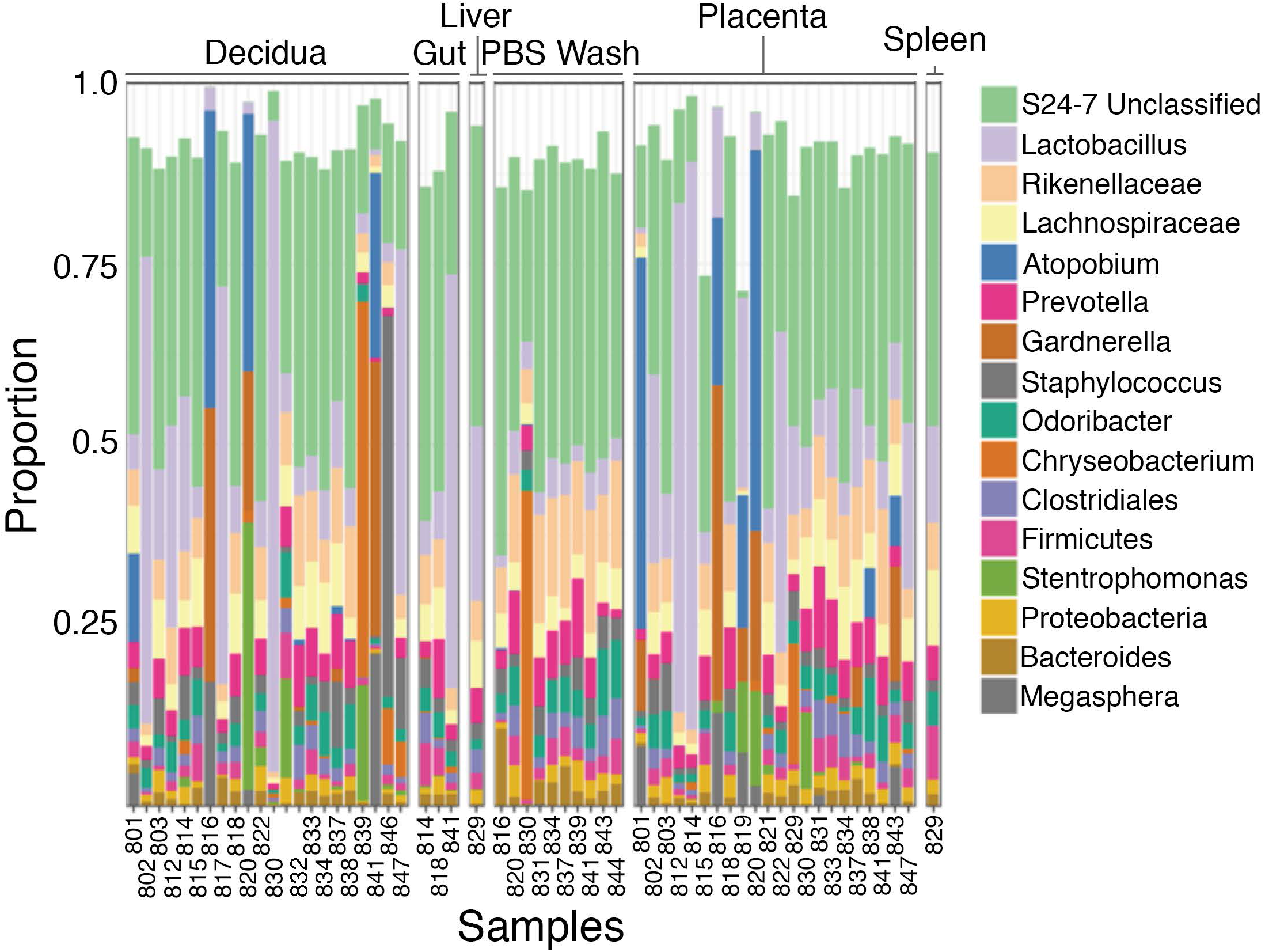
Taxonomic diversity between samples. The bacteria identified (genus) appear generally similar between decidua, placenta, and fetal tissues and phosphate buffered saline (PBS) wash fluid for most samples.

### Decidual Immune Cell Phenotyping

Resident decidual T cells are characterized by flow cytometry to investigate whether immunological responses correlate with perfusion of the placenta occurring at approximately 10 weeks gestation. T cells (defined as CD45+/CD3+ cells) are profiled for frequencies of decidual effector (CD8+) and helper (CD4+) phenotypes, frequencies of decidual Th1 (CXCR3+/CCR6-), Th2 (CXCR3-/CCR6-), and Th17 (CXCR3-/CCR6+) subsets, resident memory (CD45RO+) and activated (HLA-DRII+) T regulatory cells (Tregs; CD4+/CD25+/FoxP3+), and activated cytotoxic T cells (CD8+/HLA-DRII+). Samples are grouped into early (5 to 7 weeks gestation), mid (8-10 weeks) and late (over 10 weeks) to compare these T cell subsets during these developmental periods. The flow cytometry gating strategy is shown in Supplemental Fig. S12.

The cell type is the most significant factor driving differences in cell proportions among the groups (*p* = <.001), although an interaction between the timing of the first trimester and cell identity is also detected (*p* = .0192). Assessment of total and activated frequencies of helper and cytotoxic T cells shows that proportions do not alter across early, mid, and late time points (Fig. 5a). Similarly, Th1, Th2, Th17, and Treg frequencies do not significantly differ although Th2 frequencies showed a trend towards increasing within late decidual samples (*p* = .07; Fig. 5b and c). Frequencies of activated and memory decidual Tregs, however, do show alterations between the early, mid, and late time points. Specifically, activated Tregs increased in the mid and late time points relative to the early time point, while memory Tregs proportions showed a reciprocal decrease (Fig. 5d and e). Importantly, no significant correlation was observed between placental read counts and frequencies of activated CD4+ T cells, activated CD8+ T, memory Tregs, and the Th1/Th2 T cell ratio (Supplemental Fig. S13).

**Figure 5.**
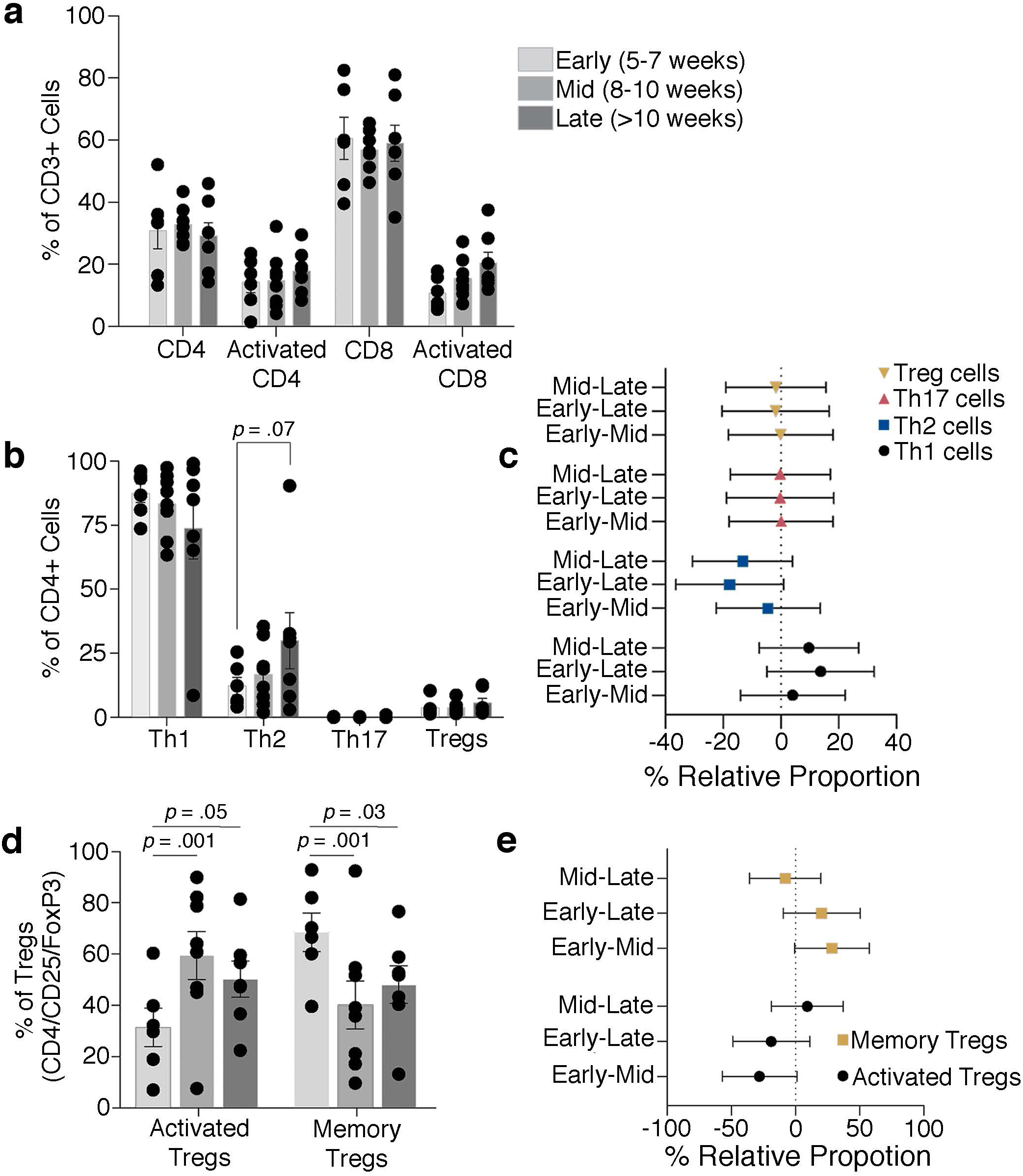
T cell dynamics through the late first to early second trimester. Proportions of total and activated decidual CD4 and CD8 T cells (a) and Th1, Th2, Th17, and Treg cells (b,c) show no significant difference through the first trimester, although there is a trend towards an increase in Th2 cells. Activated and memory Tregs levels change with increasing gestational age (d,e). Bars depict group means and error bars represent 1 standard deviation.

## Discussion

This purpose of this pilot study is to address the hypothesis that there is an increase in bacterial abundance and diversity in placental tissue and changes in resident decidual T cells after onset of maternal perfusion in the late first trimester of pregnancy. Quantitative 16S PCR of samples from the mid-first to mid-second trimester of pregnancy, which spans the pre-to post-placental perfusion timeframe, reveals detectable bacterial 16S gene signals in placental, decidual, and organ tissue samples. A weak positive correlation is found between bacterial abundance in placental tissue and gestational age in the context of no correlation for decidual samples that appears to satisfy the alternative hypothesis; however, this trend does not reach statistical significance. Moreover, comparisons of overall abundance and pre- and post-perfusion mean abundance as well as microbiome diversity show no statistically significant differences. Outliers in the taxonomic compositions of the placental, decidual, and organ tissue samples are comprised of common vaginal, gastrointestinal, and environmental bacteria suggesting contaminants introduced at the time of termination and/or sampling, although the possibility that some or all may represent nascent intrinsic placenta microbiota becoming detectable above the contamination threshold cannot be completely excluded. Together, these data argue against initiation of a placental microbiome with onset of maternal perfusion.

A shift in T regulatory cell populations in the decidua at approximately 10 weeks gestational age could be indicative of an immunological response to bacterial colonization of the placenta and provide supportive evidence for initiation of a placental microbiome with perfusion. CD4+CD25+FOXP3+ T-regulatory cells show a slight shift toward an activated memory phenotype, characterized by CD45RO+HLA-DRII+, as the first trimester progresses, which might indicate an initial reaction to establishment of placental microbiota. The proportions of CD3+ T cells, both CD4+ T helper cells and cytotoxic CD8+ cells, however, remained mostly consistent across samples from week 5 to week 13 of gestation. Moreover, no significant changes are detected in activation states of Th and cytotoxic T cells, which suggests there is no response to an immunological trigger such as the establishment of bacterial communities. The increase in HLA-DRII+ T regulatory cells is unlikely to be associated with a response to bacteria given the absence of a significant Th1 or Th2 response and is better explained by increased exposure to fetal antigens with the onset of placental perfusion.

Although examination of fetal organ tissue is not the focus of this study, organ tissue retrieved from a subset of samples and show significantly increased 16S copies compared to placenta and decidua and a moderate but statistically insignificant correlation between 16S copy abundance and gestational age. Since these tissues were intermixed with the placental and decidual samples and processed concurrently, this difference is not adequately explained by variation in contamination. This finding appears to support the concept of transfer of bacteria and/or bacterial DNA from the maternal circulation to the fetus and suggests bacteria and/or bacterial DNA may accumulate in fetal tissues even if they do not accrue in placenta; however, the number of organ tissue samples in this study is small and this would need to be properly addressed by a larger study.

There are few prior analyses of human intrauterine microbiome development in early pregnancy but there have been reports of bacteria identified in placental and fetal tissues in the first and second trimesters [4,9,13,33]. Of particular relevance is a recent study by Mishra et al. that found bacterial colonization of placental and fetal tissues between 12-22 weeks gestational age [12]. It is possible that the apparent absence of a placental microbiome in our data is due to our samples being derived at an earlier gestational age (mean 9.5 weeks), although our observation of a trend towards increased bacterial signal in placenta with increasing gestational age could be interpreted the earliest indications of a developing placental microbiome and increased bacterial abundance in organ tissues sampled between 11 and 7 weeks gestational age could represent an early point in fetal involvement. Microbiome analyses of fetal mice have produced conflicting results with some studies concluding that murine placental and fetal microbiomes develop during pregnancy [10,34,35], while others found no convincing evidence that this occurs [19,36], Of those studies promoting the presence of a murine placental microbiome, some suggest it develops later in gestation [10,35].

Selected bacterial strains of human gut microbiota can induce T-regulatory or Th17 cells [37,38]. Given the evidence that microbial communities can modulate T cell function, we inferred that the induction of a fetal microbiome concurrent with perfusion of the placenta at the end of the first trimester would lead to a shift in T cell phenotypes at the maternal-fetal interface, such as increased proportions of Th17 cells, Tregs or HLA-DRII+ cells [39]. The T cell proportions observed here are generally consistent with previous studies of T cell populations at the maternal-fetal interface [40], though we find a higher population of Th1 cells based on the CCR6-CXCR3+ expression alone compared to previous reports including additional markers and transcription factor expression [41].

This study is subject to limitations that need to be considered in the interpretation of these findings, the most important of which is the effect of bacterial contamination on microbiome analysis. We structured our study to assess for relative, rather than absolute, changes in bacterial abundance and diversity based on data suggesting that contamination is relatively consistent in abundance and diversity between samples [15,20]. In this respect we endeavored to minimize variability in contamination between samples by batching handling of specimens, standardizing sampling and testing protocols, and using the same equipment and operators. We expect changes over background contamination to be detectable in comparison to decidua; however, the sensitivity of this approach in early pregnancy is unclear and variance in contaminants between samples could overwhelm real changes. As such, we cannot exclude the possibility that a placental microbiome begins to develop in early pregnancy, only that it is not detectable above the level of background sample contamination in this study. Factors such as DNA extraction, the16S rRNA variable region amplified, and differential amplification between samples, which has a greater proportional effect in low biomass samples, may have contributed to masking more subtle changes [5,15], although differences in these effects between samples should be negligible.

Implicit to our study is the assumption, based on previous placental microbiome studies, that hematogenous spread to the placenta is the primary source of bacteria. It is possible that other routes (transcervical, transmembrane) are involved, which this study is not designed to address and if significant could mask pre- and post-perfusion differences. Our placental samples are derived from chorionic villus tissue, which may not be fully representative of the entire placental microbiome [21]; however, placental villi are the interface between the placenta and maternal circulation and would be the most representative of bacteria introduced by this route. It is important to acknowledge that we employ the presence of bacterial DNA and immunological reaction as markers for the presence of bacteria, but this does not equal the presence of intact, viable bacteria. In this respect, it is notable that bacteria were not histological apparent in representative tissue from each sample. Regarding flow cytometry, it is possible that the assays employed were not sensitive enough to detect subtle shifts in decidual T cell behavior. Investigating the transcriptional profiles of resident T cells and their TCR repertoire via sequencing could elucidate more nuanced roles for decidual T cells, perhaps better revealing their affinity for commensal bacteria or alternative roles to their counterparts in circulation.

The purpose of this pilot study is to assess for changes in the abundance and/or diversity of bacteria in placental tissue after initiation of perfusion during early human pregnancy using microbiome and flow cytometric analyses as indirect markers. The data argue against development of the placental microbiome in early pregnancy, although it is possible that it is present but below the limit of detection during the timeframe of this study. Additional investigations in larger cohorts are warranted to further explore this possible association.

## Acknowledgments

The authors thank the study participants for their generous cooperation, the associated medical staff and BC Children’s Hospital Biobank for their assistance and Dr. David Goldfarb for his suggestions on study design and manuscript preparation. This study is supported by a BC Children’s Hospital Foundation Fetal Growth and Development Catalyst Grant.

