## Supplementary Figures and Tables for "Assessment of the Human Placental Microbiome in Early Pregnancy"

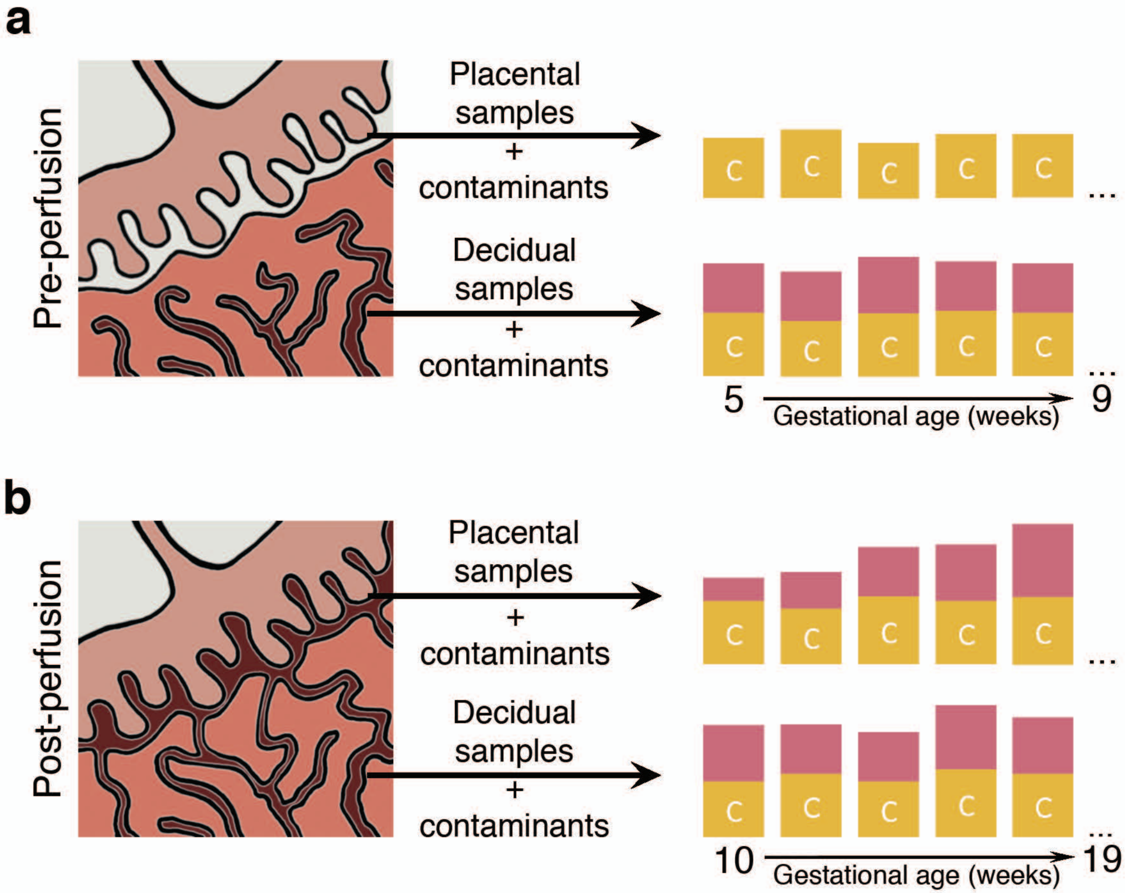


Supplemental Figure S1. The microbiome component of this study is designed to detect an increase in placental bacterial DNA compared to decidua above background bacterial contamination. During the anoxic phase of embryonic development the placenta is not perfused by the maternal circulation and not exposed to blood-borne bacteria. During this pre-perfusion phase,only bacterial contaminants introduced during delivery sampling (yellow boxes, labeled "C") should be detectable in placental samples while decidual samples contain bacteria derived from the maternal circulation (red boxes) as well as contaminants (a). After onset of perfusion the placenta is exposed to blood-borne bacteria that should become detectable as increasing bacterial abundance and/or diversity over time compared to decidua (b).

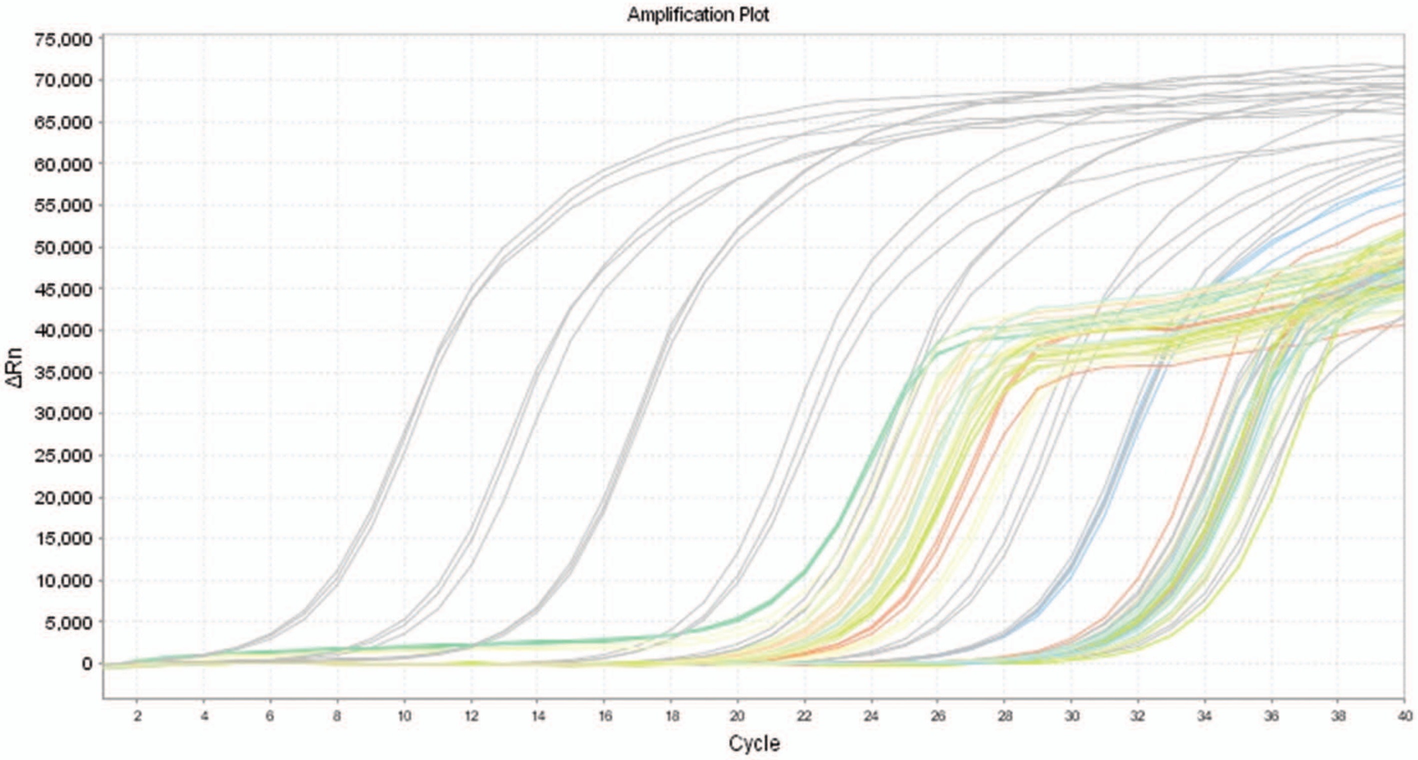


Supplemental Figure S2. Quantitative 16S PCR amplifications curves.The left cluster of colored curves represent the tissue samples and the right cluster of colored curves represent the PBS wash samples. The grey curves represent 10 fold dilutions of the bacterial 16S rRNA control.

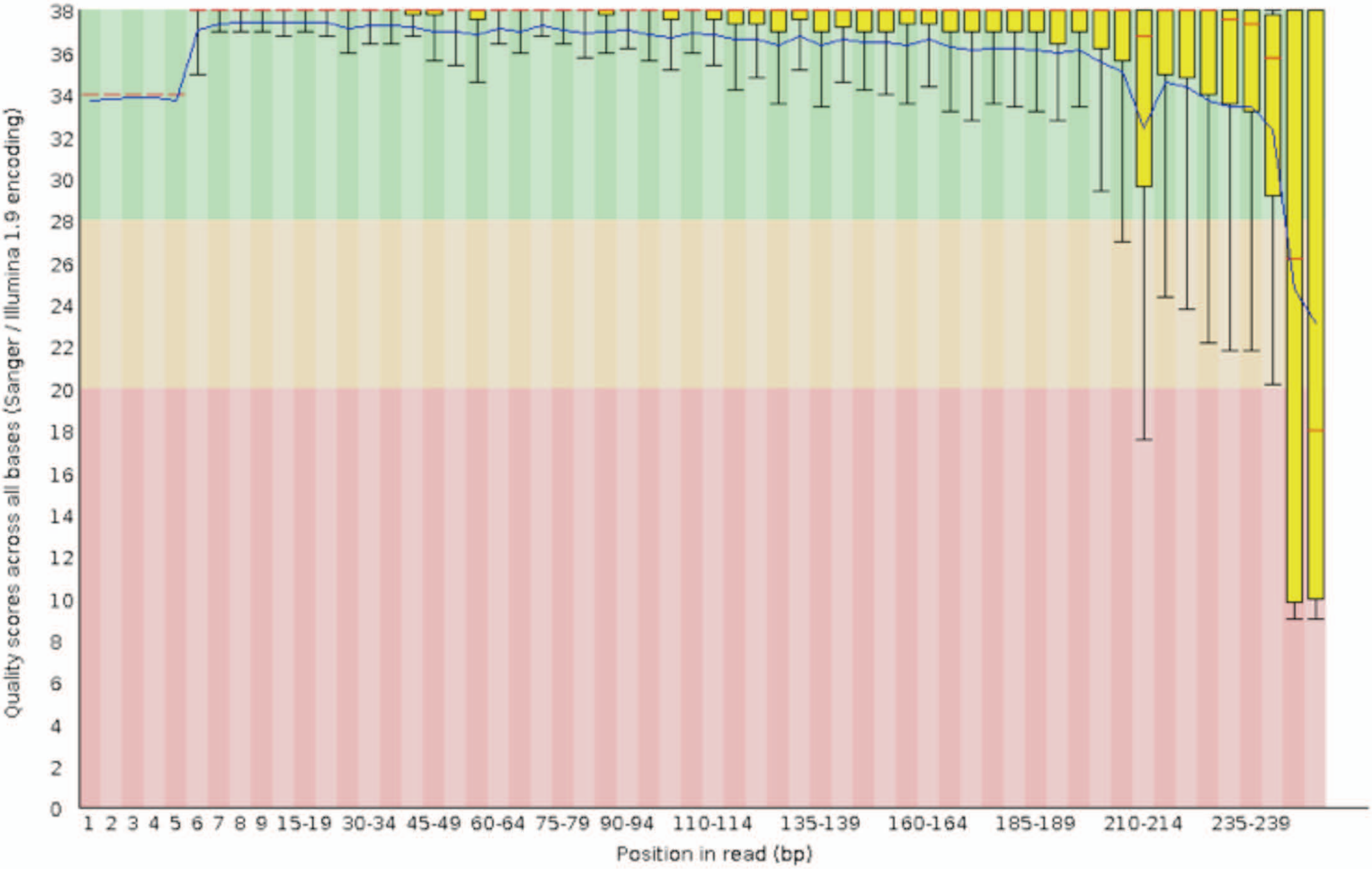


Supplemental Figure S3. Per-base raw Phred33 scores of the forward sequencing reads. Scores are determined using FastQC 0.11.5 and assess the accuracy of sequencing and indicates the probability that a particular base has been called incorrectly Green (Q score 28 or greater, probability of an incorrect call 1/933 or less) indicates high quality, yellow reasonable, and red poor. The blue line indicates mean quality, the red line indicates median value, yellow box the 25-75th percentile, and the whiskers the 10-90th percentile.

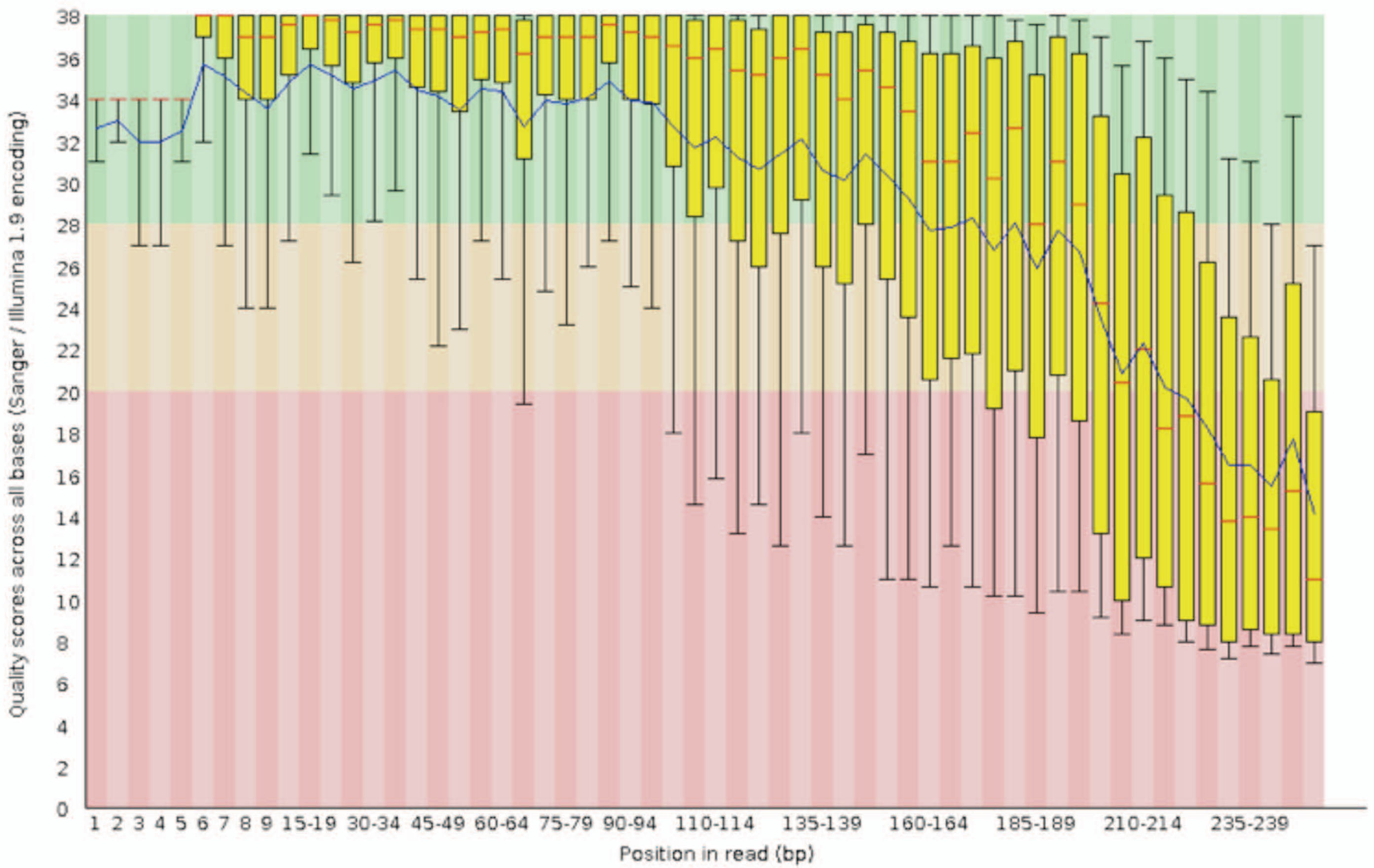


Supplemental Figure S4. Per-base raw Phred33 scores of the reverse sequencing reads. Scores are determined using FastQC 0.11.5 and assess the accuracy of sequencing and indicates the probability that a particular base has been called incorrectly. Green (Q score 28 or greater, probability of an incorrect call 1/933 or less) indicates high quality, yellow reasonable, and red poor. The blue line indicates mean quality, the red line indicates median value, yellow box the 25-75th percentile, and the whiskers the 10-90th percentile.

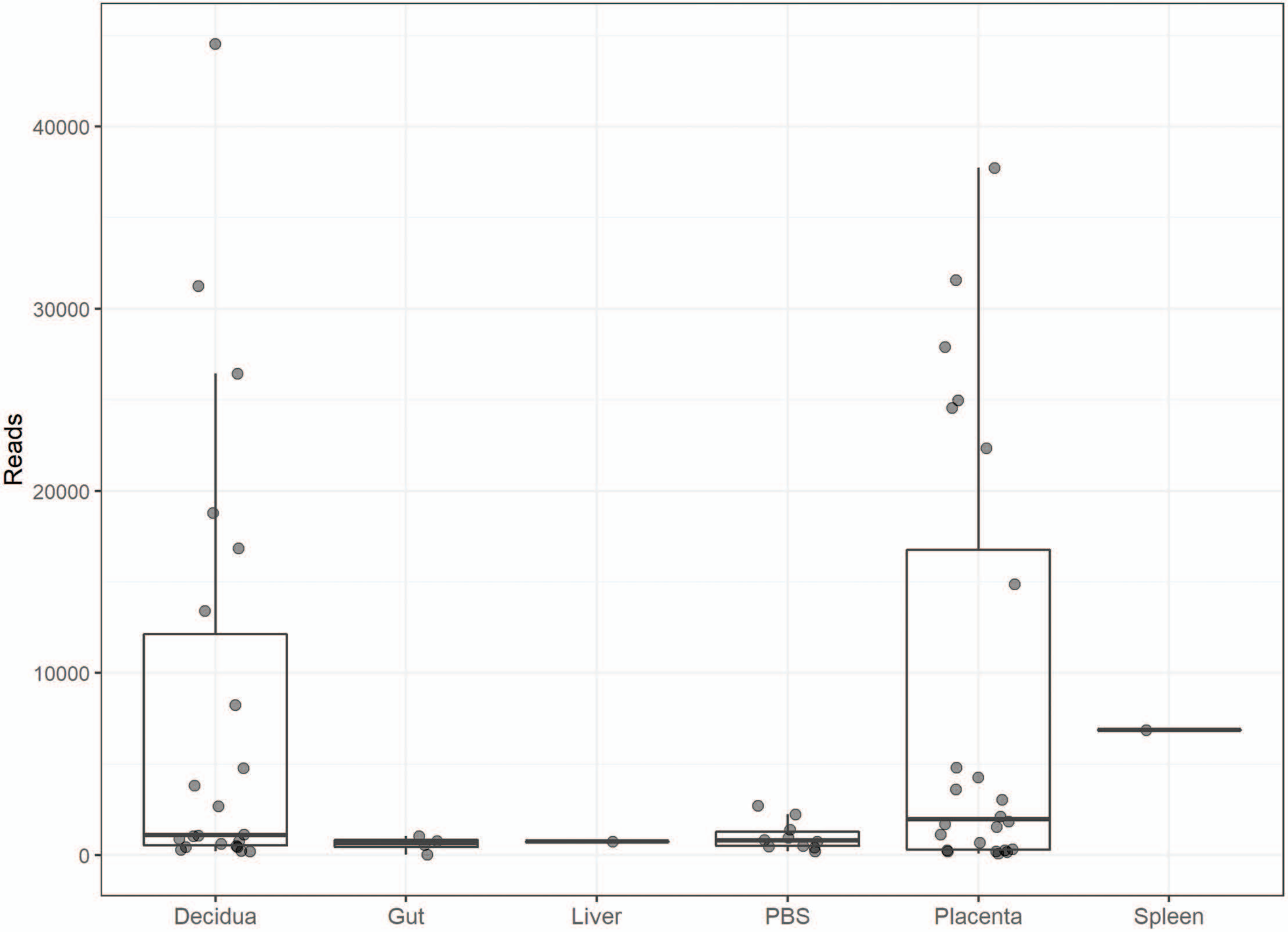


Supplemental Figure S5. Total number of quality filtered reads per sample. Reads include high-quality sequences that align with 16Sv4, are clustered into OTUs, and are assigned taxonomic classification. Ambiguous or low-quality data are excluded from subsequent analyses.

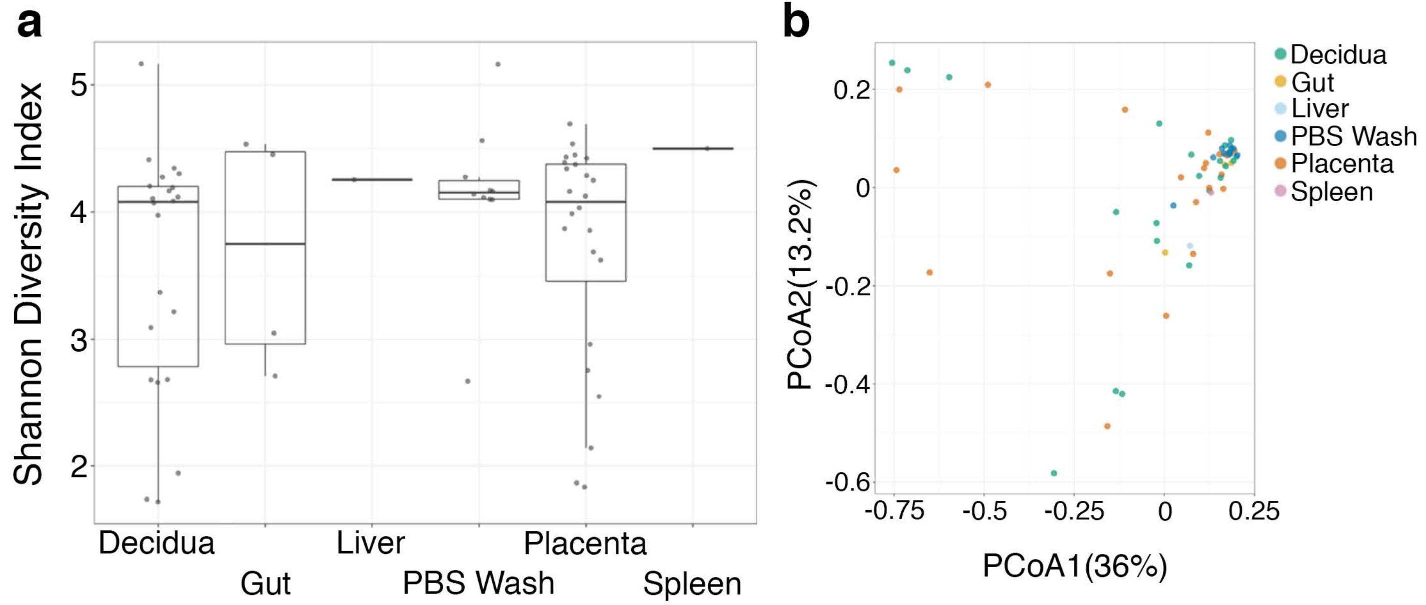


Supplemental Figure S6. Bacterial diversity between samples. Average diversity is similar between maternal decidua, placenta, fetal tissue, and phosphate buffered saline (PBS) wash samples (a). Principal coordinate analysis demonstrating similarity between samples (b). Box plots encompass the mean, upper and lower quartiles.

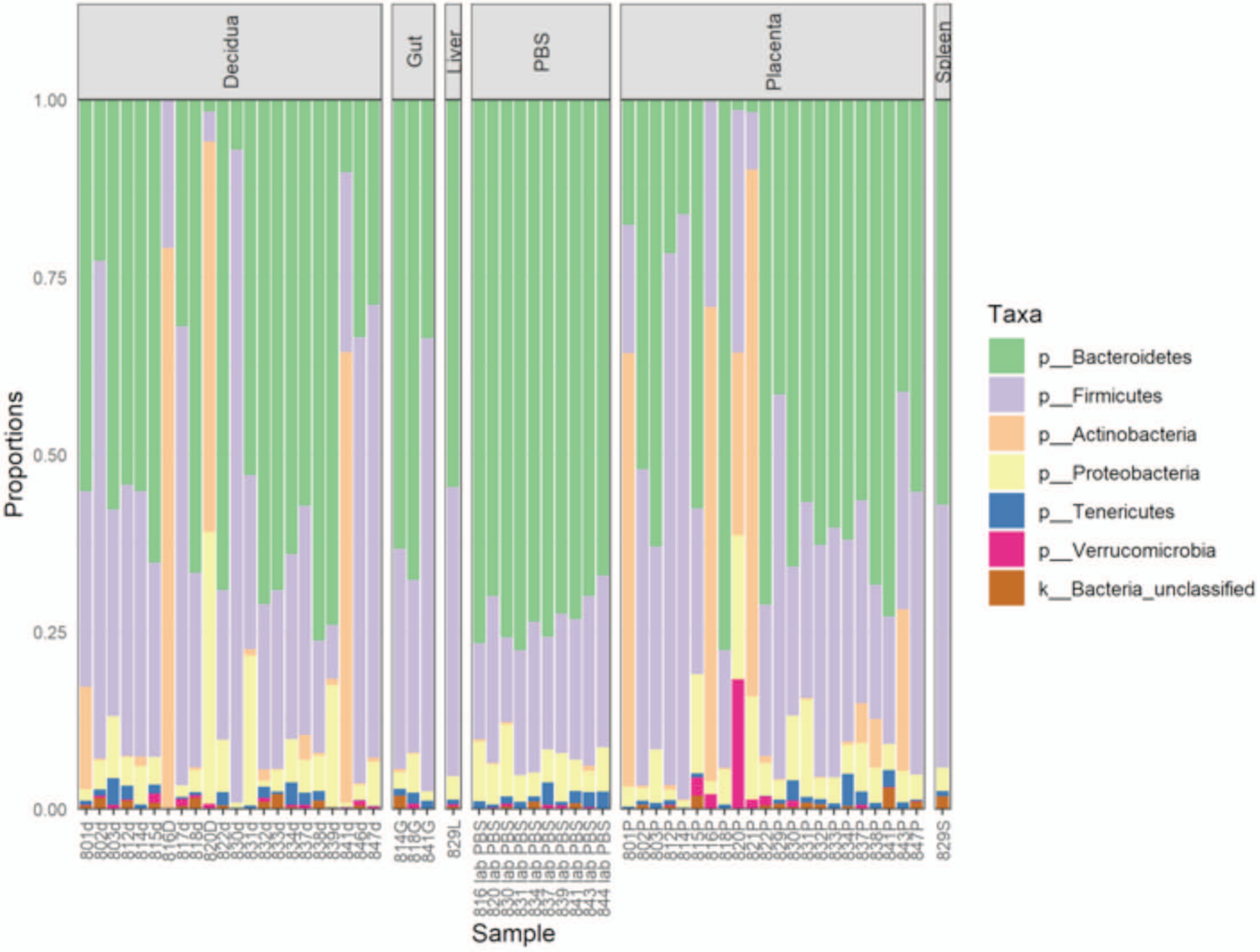


Supplemental Figure S7. Taxonomic distribution by phylum. Decidua, placenta, and fetal tissues and phosphate buffered saline (PBS) wash samples. Unfilled portion of the bar plots represent lower abundance taxa.

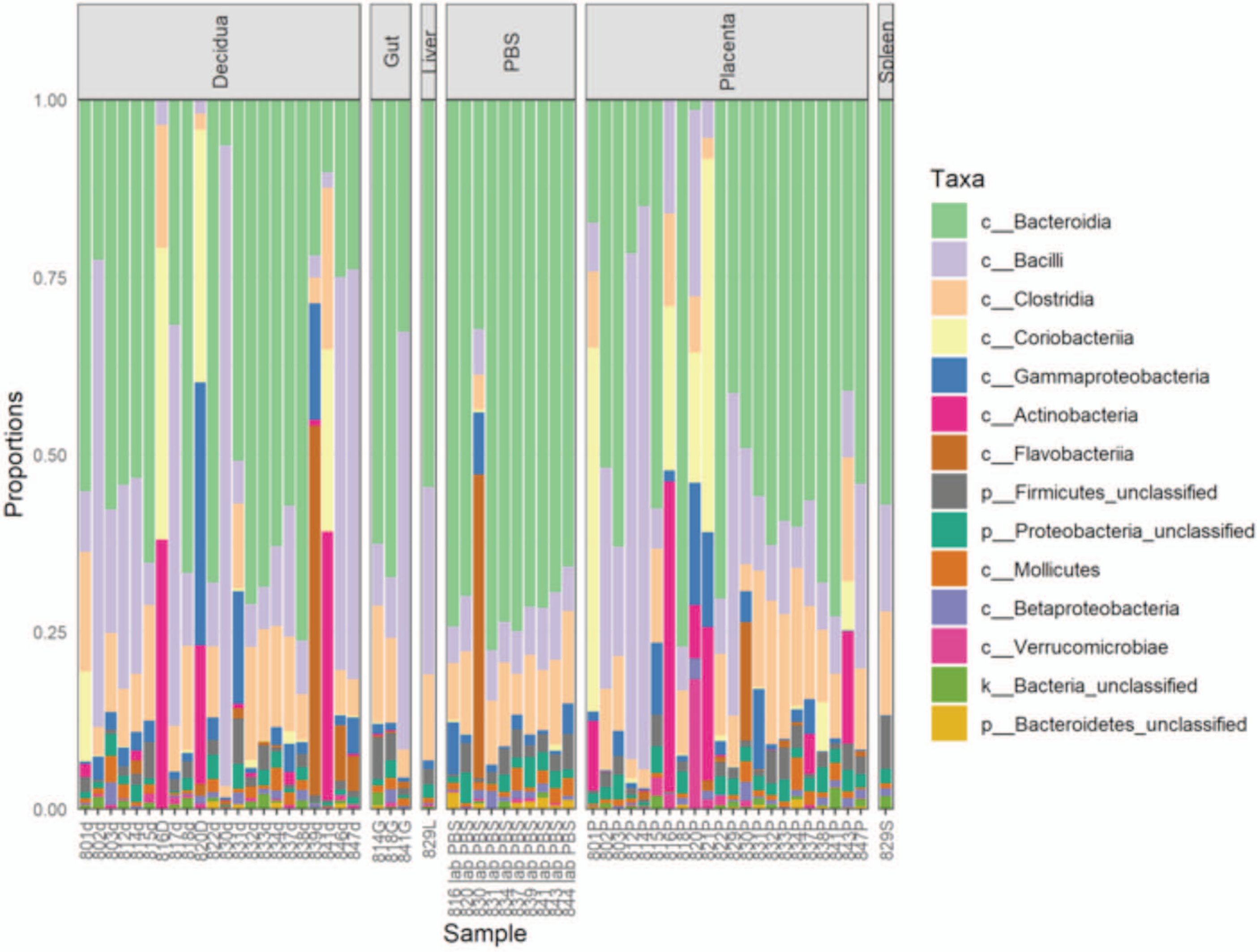


Supplemental Figure S8. Taxonomic distribution by class. Decidua, placenta, and fetal tissues and phosphate buffered saline (PBS) wash samples.

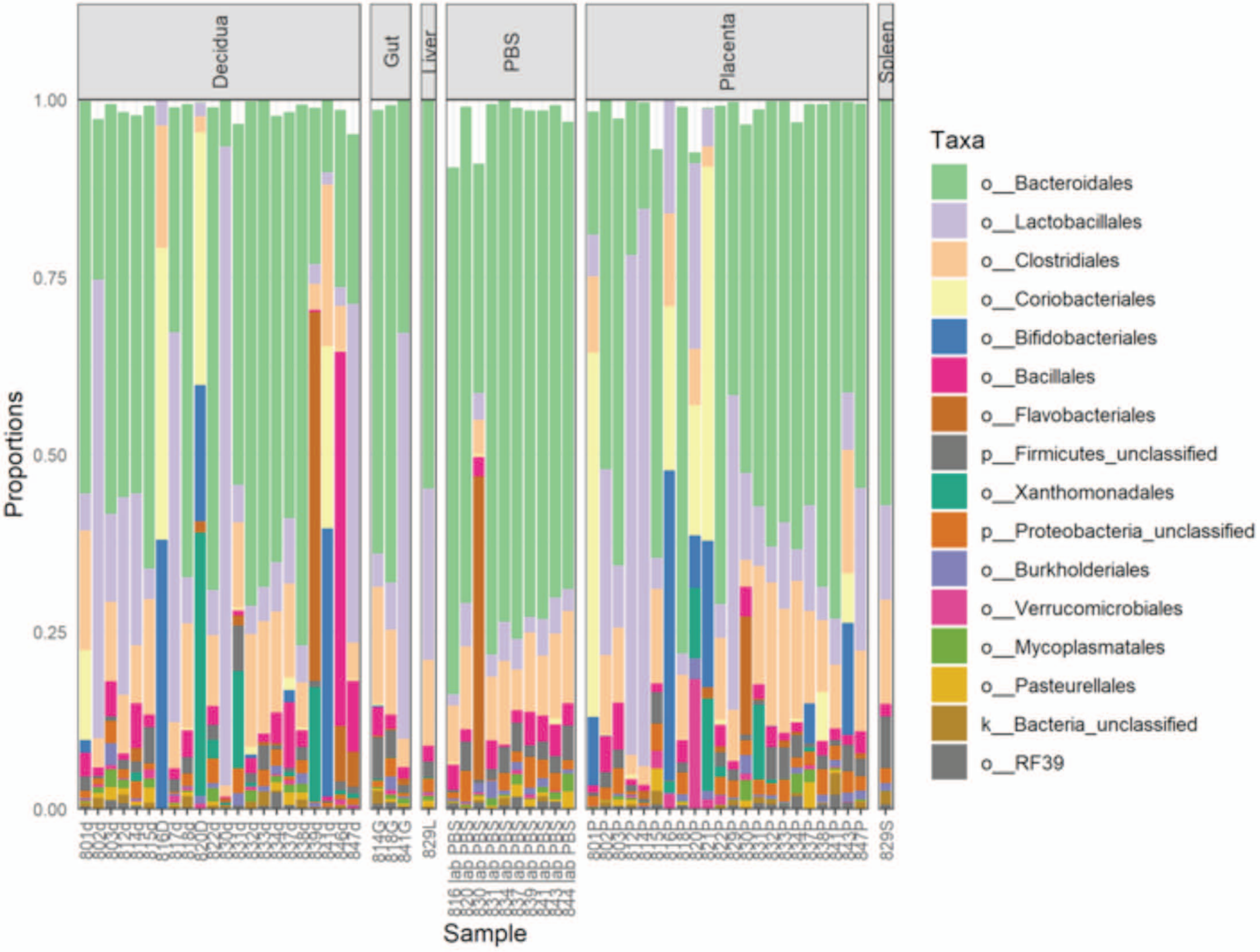


Supplemental Figure S9. Taxonomic distribution by order. Decidua, placenta, and fetal tissues and phosphate buffered saline (PBS) wash samples. Unfilled portion of the bar plots represent lower abundance taxa.

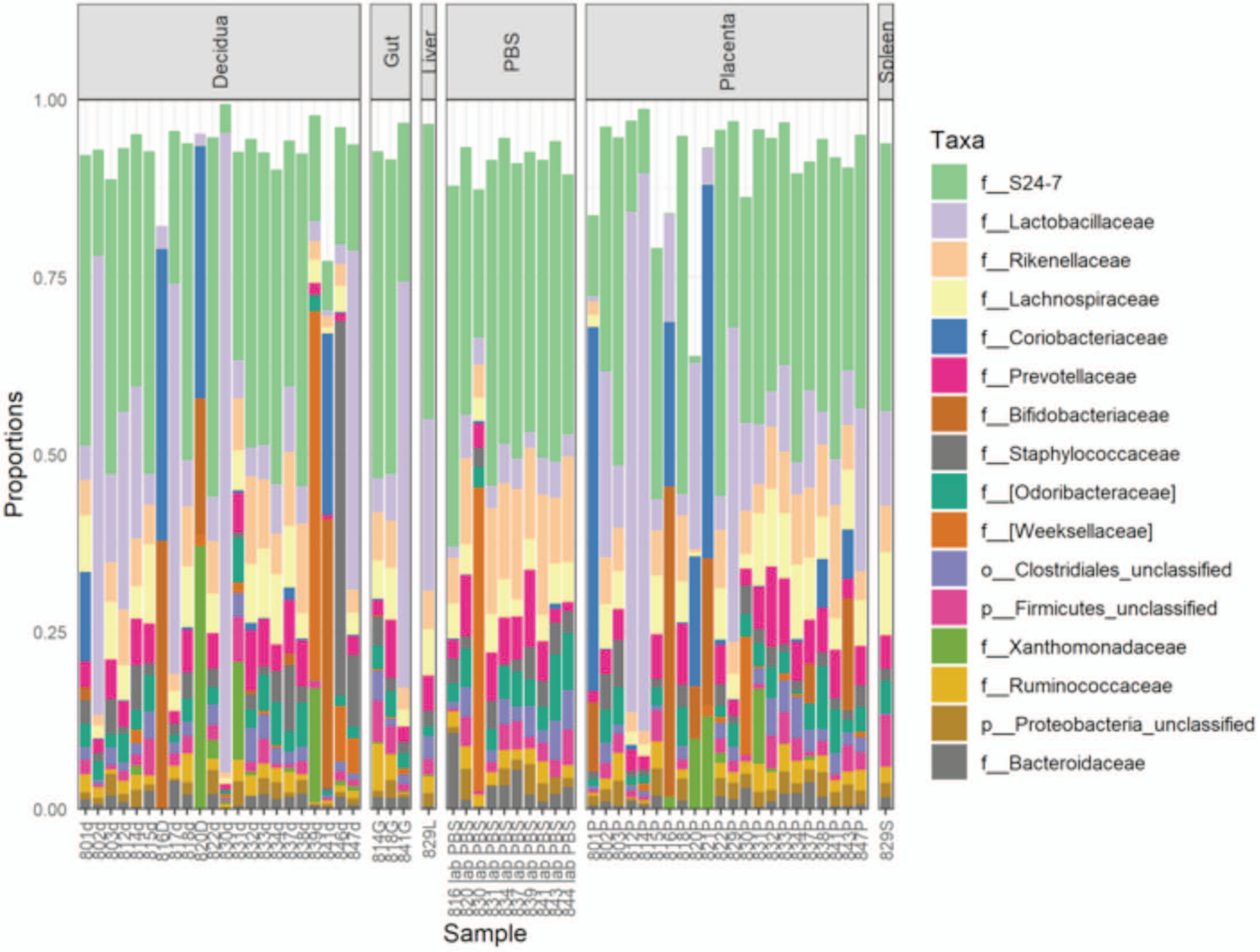


Supplementarl Figure S10. Taxonomic distribution by family. Decidua, placenta, and fetal tissues and phosphate buffered saline (PBS) wash samples. Unfilled portion of the bar plots represent lower abundance taxa.

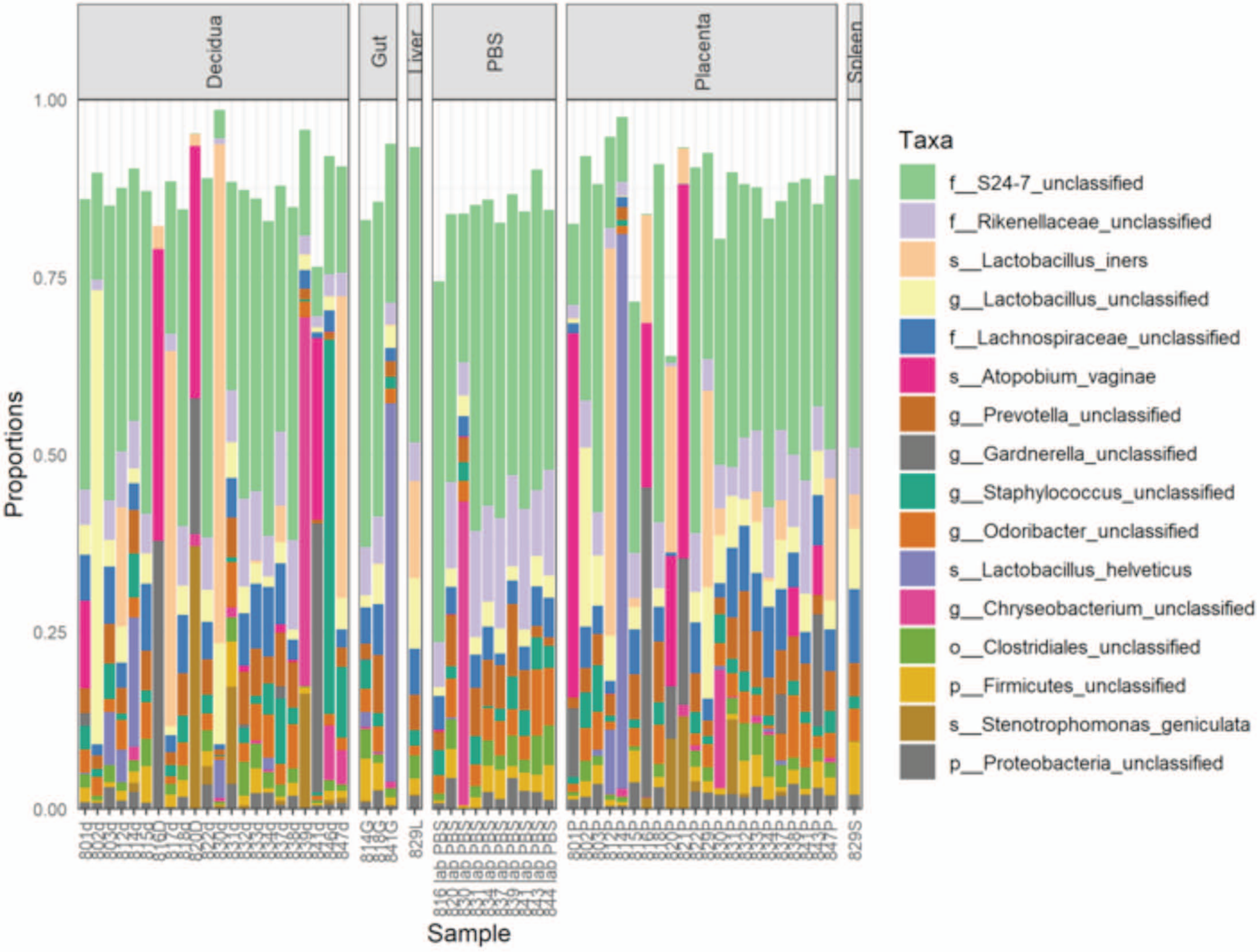


Supplemental Figure S11. Taxonomic distribution by genus/species. Decidua, placenta, and fetal tissues and phosphate buffered saline (PBS) wash samples. Unfilled portion of the bar plots represent lower abundance taxa.

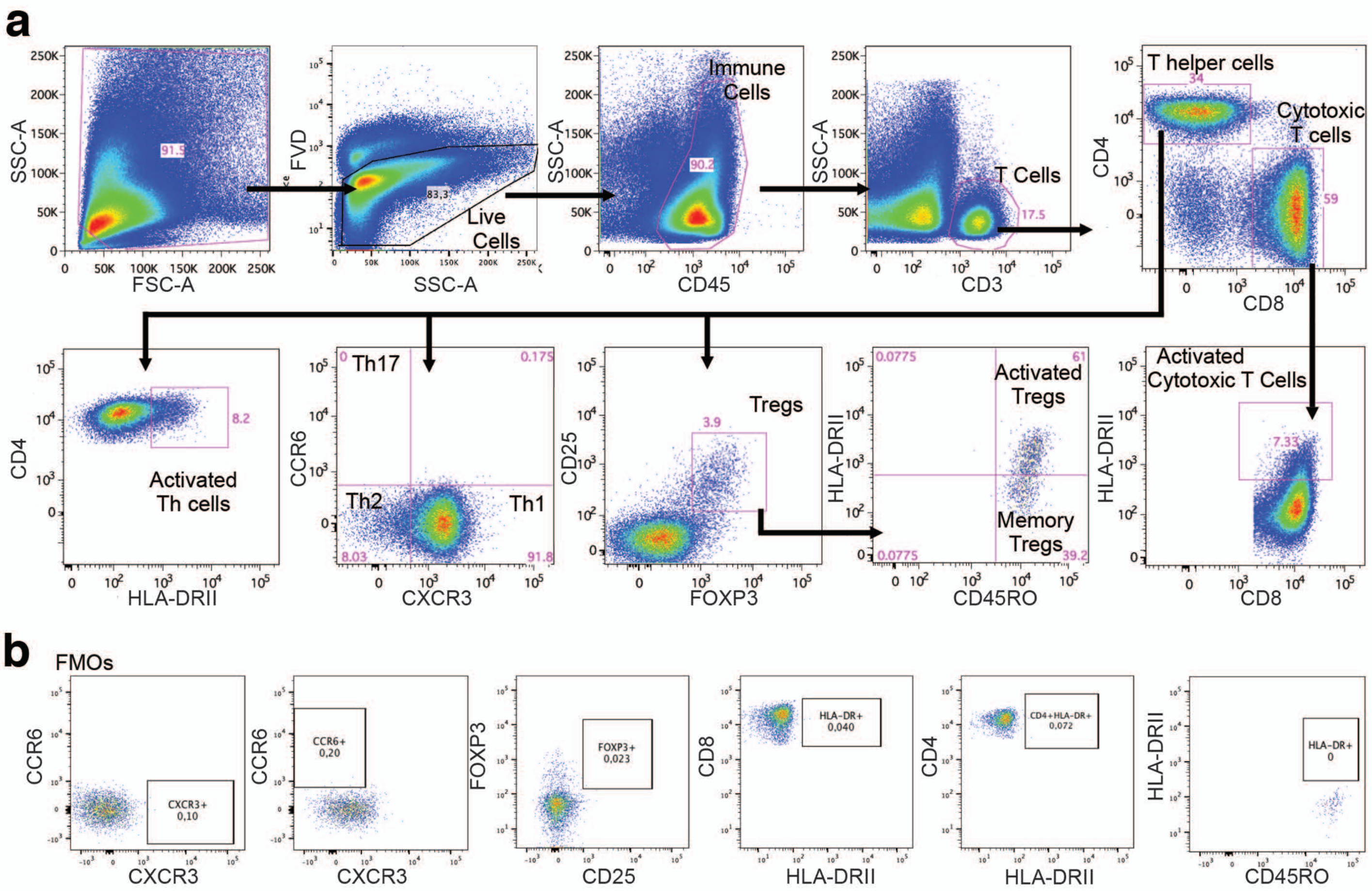


Supplemental Figure S12. Flow cytometry gating strategies. T cell subsets (a) and fluorescence minus one (FMO) controls (b).

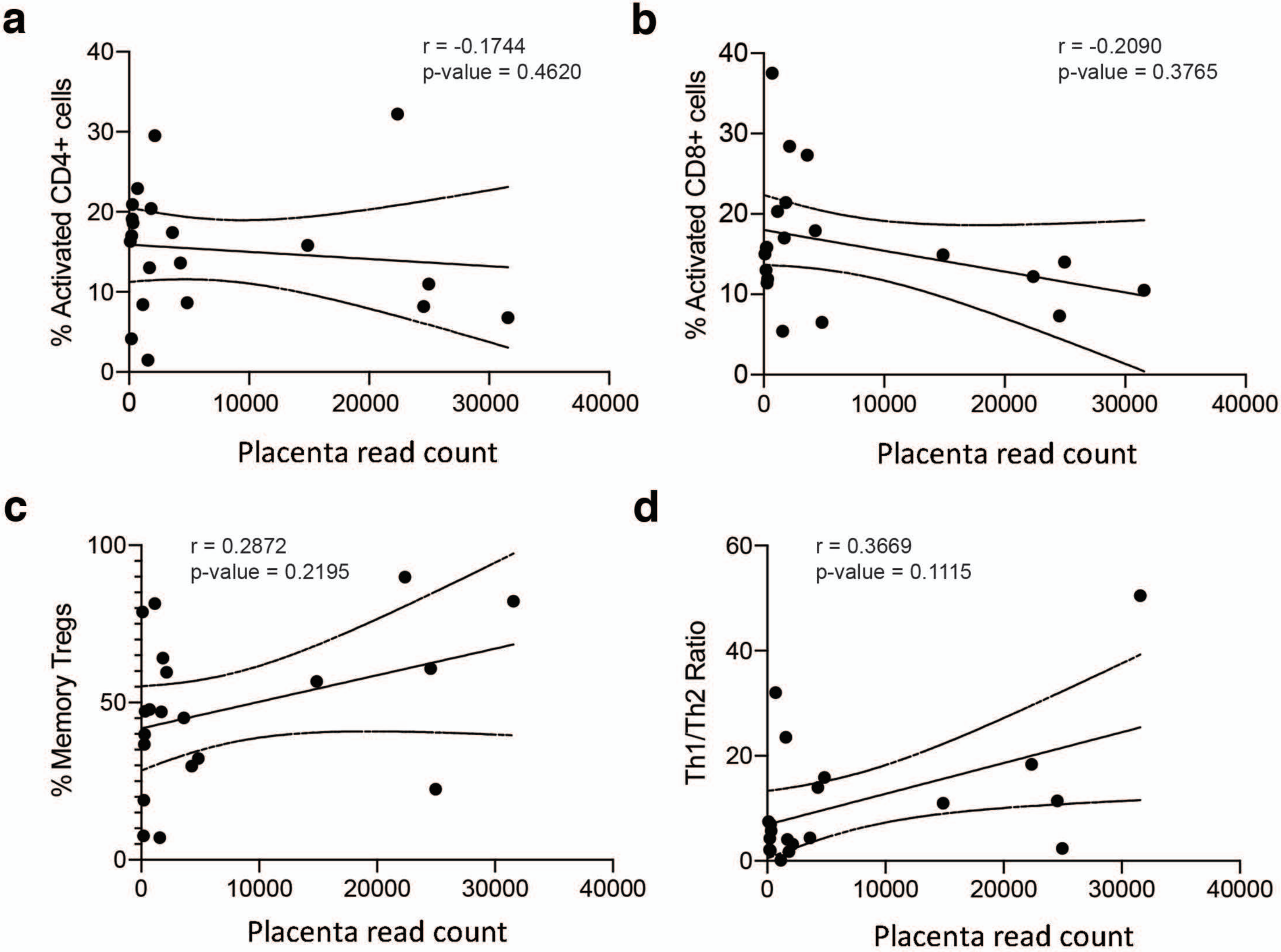


Supplemental Figure S13. Correlation between placental read counts and frequencies of activated CD4+ T cells (a), activated CD8+ T cells (b), memory Treg cells (c), and the Th1/Th2 ratio (d).

| Sample # | Age | BMI | GA | Tissues | Question Answers |
|---|---|---|---|---|---|
| 801 | 35 | 21.3 | 5.6 | Placenta, decidua, decidual immune cells | Q1: No. Q2: No. Q3: No. Q4: No. Q5: No. |
| 802 | 40 | 32.8 | 9.1 | Placenta, decidua | Q1: No. Q2: No. Q3: No. Q4: No. Q5: No. |
| 803 | 21 | 26 | 7.4 | Placenta, decidua, decidual immune cells | Q1: No. Q2: No. Q3: No. Q4: No. Q5: No. |
| 812 | 22 | 18.1 | 5.3 | Placenta, decidua, decidual immune cells | Q1: No. Q2: No. Q3: No. Q4: No. Q5: No. |
| 814 | 21 | 22.7 | 11 | Placenta, decidua, fetal bowel | Q1: Yes. Q2: Yes. Q3: No. Q4: Yes. Q5: No. |
| 815 | 21 | 20.1 | 8.5 | Placenta, decidua, decidual immune cells | Q1: No. Q2: No. Q3: No. Q4: No. Q5: No. |
| 816 | 40 | 22.6 | 9.3 | Placenta, decidua, decidual immune cells | Q1: No. Q2: No. Q3: No. Q4: No. Q5: No. |
| 817 | 26 | 21.5 | 6.6 | Placenta, decidua, decidual immune cells | Q1: No. Q2: Yes. Q3: No. Q4: Yes. Q5: No. |

| | | | | | |
|---|---|---|---|---|---|
| 818 | 39 | 28.3 | 13.6 | Placenta, decidua, decidual immune cells, fetal bowel | Q1: No. Q2: No. Q3: No. Q4: No. Q5: No. |
| 820 | 27 | 36.5 | 10 | Placenta, decidua, decidual immune cells | Q1: No. Q2: No. Q3: No. Q4: No. Q5: No. |
| 821 | 27 | 23.9 | 8.1 | Placenta, decidua, decidual immune cells | Q1: No. Q2: Yes. Q3: No. Q4: Yes. Q5: No. |
| 822 | 32 | 20.4 | 8.3 | Placenta, decidua, decidual immune cells | Q1: No. Q2: No. Q3: No. Q4: No. Q5: No. |
| 829 | 30 | 31.1 | 17.1 | Placenta, Fetal gut, liver, spleen | Q1: No. Q2: Yes. Q3: No. Q4: No. Q5: No. |
| 830 | 23 | 38.8 | 10 | Placenta, decidua, decidual immune cells | Q1: No. Q2: No. Q3: No. Q4: No. Q5: No. |
| 831 | 24 | 22.3 | 8.6 | Placenta, decidua, decidual immune cells | Q1: No. Q2: Yes. Q3: No. Q4: No. Q5: No. |
| 832 | 29 | 27.5 | 9.1 | Placenta, decidua, decidual immune cells | Q1: No. Q2: No. Q3: No. Q4: Yes. Q5: No. |

| | | | | | |
|---|---|---|---|---|---|
| 833 | 30 | 22.1 | 7.5 | Placenta, decidua, decidual immune cells | Q1: No. Q2: No. Q3: No. Q4: No. Q5: No. |
| 834 | 34 | 34.4 | 11.5 | Placenta, decidua, decidual immune cells | Q1: No. Q2: Yes. Q3: No. Q4: No. Q5: No. |
| 837 | 30 | 23 | 9.3 | Placenta, decidua, decidual immune cells | Q1: No. Q2: No. Q3: No. Q4: No. Q5:No. |
| 838 | 27 | 26.5 | 10 | Placenta, decidua, decidual immune cells | Q1: No. Q2: No. Q3: No. Q4: No. Q5: No. |
| 839 | 23 | 29.1 | 7.3 | Placenta, decidua, decidual immune cells, | Q1: No. Q2: No. Q3: No. Q4: No. Q5: No. |
| 841 | 21 | 22.7 | 14.4 | Placenta, decidua, decidual immune cells, fetal gut | Q1: No. Q2: No. Q3: No. Q4: No. Q5: No. |
| 843 | 19 | 30.8 | 12 | Placenta, decidua, decidual immune cells | Q1: No. Q2: Yes. Q3: No. Q4: No. Q5: No. |
| 846 | 39 | 33.3 | 9.6 | Placenta, decidua, decidual immune cells | Q1: No. Q2: No. Q3: No. Q4: No. Q5: No. |
| 847 | 27 | 22.6 | 9.5 | Placenta, decidua, decidual immune cells | Q1: No. Q2: Yes. Q3: No. Q4: No. Q5: No. |

Supplementary Table S1. Patient cohort demographics and answers to questions 1-5. Q1: “Did you take anti-hypertensive medications pre/post pregnancy?” Q2: “Did you smoke?” Q3: “Are you a diabetic/do you have diabetes?” Q4: “Do you take anti-inflammatory medications?”Q5: “Are you presently taking antibiotics?”.

| Sample | Mean CT | Quantity | Undiluted Quantity |
|---|---|---|---|
| 801P | 23.82 | 2881 | 28806 |
| 801D | 23.62 | 3250 | 32500 |
| 802P | 22.63 | 6024 | 60242 |
| 802D | 23.51 | 3492 | 34922 |
| 803P | 23.63 | 3231 | 32306 |
| 803D | 22.89 | 5107 | 51068 |
| 812P | 22.64 | 5987 | 59866 |
| 812D | 22.04 | 8656 | 86556 |
| 814P | 24.13 | 2365 | 23651 |
| 814D | 24.12 | 2382 | 23821 |
| 814G | 21.75 | 10418 | 104184 |
| 815P | 21.31 | 13653 | 136535 |
| 815D | 21.68 | 10872 | 108720 |
| 816D | 21.00 | 15000 | 15000 |
| 816P | 20.00 | 25000 | 16000 |
| 817D | 21.62 | 11246 | 112455 |
| 818P | 21.24 | 14246 | 142464 |
| 818D | 22.18 | 7962 | 79616 |
| 818G | 18.53 | 76424 | 764236 |
| 820P | 21.00 | 23000 | 5800 |
| 820D | 22.00 | 12000 | 4200 |
| 821P | 20.00 | 25000 | 8600 |
| 822P | 22.64 | 5973 | 59725 |
| 822D | 22.77 | 5532 | 55323 |
| 829P | 23.01 | 4744 | 47435 |
| 829G | 17.62 | 134640 | 1346397 |
| 829S | 22.49 | 6582 | 65816 |
| 829L | 18.24 | 91565 | 915650 |
| 830P | 21.74 | 10485 | 104855 |
| 830D | 23.14 | 4386 | 43858 |
| 831P | 25.60 | 950 | 9497 |
| 831D | 23.62 | 3252 | 32520 |
| 832P | 24.09 | 2425 | 24251 |
| 832D | 24.38 | 2028 | 20282 |
| 833P | 22.58 | 6201 | 62005 |
| 833D | 24.28 | 2165 | 21647 |
| 834P | 22.61 | 60767 | 60767 |
| 834D | 20.82 | 18560 | 185600 |
| 837P | 22.52 | 6434 | 64342 |
| 837D | 22.55 | 6304 | 63042 |
| 838P | 26.96 | 409 | 4087 |

| | | | |
|---|---|---|---|
| 838D | 27.44 | 304 | 3040 |
| 839P | 23.36 | 3828 | 38276 |
| 839D | 25.72 | 885 | 8846 |
| 841P | 31.49 | 25 | 247 |
| 841D | 24.17 | 2320 | 23202 |
| 841G | 24.02 | 2544 | 25444 |
| 843P | 24.23 | 2227 | 22266 |
| 846P | 23.17 | 4315 | 43151 |
| 846D | 24.99 | 1394 | 13938 |
| 847P | 22.74 | 5613 | 56134 |
| 847D | 21.09 | 15673 | 156732 |
| 830 PBS wash | 29.34 | 94 | 936 |
| 820 PBS wash | 32.57 | 13 | 126 |
| 834 PBS wash | 32.80 | 11 | 109 |
| 839 PBS wash | 32.86 | 10 | 105 |
| 844 PBS wash | 33.30 | 8 | 80 |
| 837 PBS wash | 33.07 | 9 | 93 |
| 841 PBS wash | 32.22 | 16 | 157 |
| 843 PBS wash | 32.83 | 11 | 107 |
| 831 PBS wash | 32.90 | 10 | 103 |
| 816 PBS wash | 32.67 | 12 | 118 |

Supplementary Table S2. Quantitative 16S PCR mean CT values and quantity by sample. P = placenta, D = decidua, G = gut (intestine), S = spleen, L = Liver, PBS = phosphate buffered saline.

| Cell Marker | Fluorophore | Supplier | Reference Number | Clone | Dilution |
|---|---|---|---|---|---|
| CD45 | PeCy5 | eBioscience | 15-0459-42 | HI30 | 1:75 |
| CD3 | PerCP710 | eBioscience | 46-0036-42 | SK7 | 1:75 |
| CD8 | APCH7 | BD Biosciences | 560179 | SK1 | 1:30 |
| CD4 | Alexa700 | BD Biosciences | 557922 | RPA-T4 | 1:50 |
| CXCR3 | APC | BD Biosciences | 561732 | 1C6/CXCR3 | 1:25 |
| CCR6 | PeCy7 | eBioscience | 25-1969-42 | R6H1 | 1:50 |
| CD25 | ef450 | eBioscience | 48-0259-42 | BC96 | 1:25 |
| CD45RO | PE | eBioscience | 12-0457-42 | UCHL1 | 1:50 |
| HLA-DR | V500 | BD Biosciences | 561225 | G46-6 | 1:50 |
| Viability Dye | FVD520 | eBioscience | 65-0867-14 | N/A | 1:1000 |
| FOXP3 | PE-CF594 | BD Biosciences | 562421 | 259D/C7 | 1:25 |

Supplementary Table S3. Flow cytometry antibodies used in this study.
